# Mapping of suitable release areas for the parasitoid *Dolichogenidea gelechiidivoris* (Marsh) for the classical biocontrol of *Tuta absoluta* (Meyrick) using temperature-dependent phenology models

**DOI:** 10.1101/2024.12.17.628750

**Authors:** Norma Mujica, Pablo Carhuapoma, Jürgen Kroschel, Jan Kreuze

## Abstract

The South American tomato leafminer, *Tuta absoluta* (Meyrick) is a major invasive pest of tomato (*Lycopersicum esculentum* Mill.) worldwide. Among the different integrated pest management strategies, biological control is most promising. *Dolichogenidea gelechiidivoris* (Marsh) is a larval endoparasitoid native to the Neotropics, where it is the dominant biological agent of *T. absoluta* along the Peruvian coast. The determination of the parasitoid’s temperature-dependent development is crucial for better predicting the potential of the parasitoid to establish in new regions and to control the target pest. Therefore, the effect of temperature on the development and reproduction of *D. gelechiidivoris* was studied at five constant temperatures ranging from 10 to 30°C in its main host *T. absoluta*. The Insect Life Cycle Modeling (ILCYM) software was used to fit nonlinear equations to collected life table data and to establish an overall phenology model to simulate life table parameters based on temperature. The parasitoid completed its life cycle at constant temperatures from 15 to 30°C; the temperature of 10°C was lethal to pupae. The theoretical lower threshold temperatures for the development of egg-larvae and pupae were 7.6°C and 10.9°C respectively. The egg-larval and pupae stages had the lowest mortality between the temperature range of 20-30°C. The lowest senescence rates for females and males were observed within the temperature range of 10–20°C. Oviposition time decreased significantly with increasing temperature from 16.7 days (10°C) to 1.6 days (35°C). Mean fecundity was highest at 20°C (74.4 eggs/female). Maximum population growth is expected around 24.3°C with a finite rate of increase, λ of 1.1088, which corresponds to a population doubling time of 6.7 days. The highest values for gross reproduction rate (GRR) and net reproduction rate (R_0_) were found between 20 and 21°C, and the shortest mean generation time (T) was observed at 30°C (19.9 d). Suitable release areas with a very high probability of establishment and potentially good control efficacy of the parasitoid are tropical and subtropical regions (e.g., countries in Southern Europe; Spain, Portugal). The potential use of the parasitoid in the context of classical biological control of *T. absoluta* is discussed.

## Introduction

Invasive species in the form of plants, animals, insects, and diseases are the largest impediment to global food security and agricultural production (FAO, 2008). Hence, maintaining food security in part requires reducing losses from endemic and invasive pests, a problem which is projected to exacerbate under climate change as temperature is one of the most important factors affecting insect pests (Kroschel, Mujica, Carhuapoma & Sporleder, 2016; Bale et al., 2002). The South American tomato leafminer, *Tuta absoluta* (Meyrick) (Lepidoptera, Gelechiidae), widespread in all countries of South America, is believed to be native to Peru where it was first described in 1917 (Desneux et al., 2010). It still has a limited distribution in Central America (Panama, and Cost Rica), but after its transatlantic invasion and first detection in Spain in 2006, the pest rapidly spread across southern Europe, North Africa, and the Near East (Urbaneja, Vercher, Navarro, Garcia Mari & Porcuna, 2007; Potting, 2009; Campos, Biondi, Adiga, Guedes & Desneux, 2017) and continued invading more than 90 countries outside of South America in Europe, Africa, and Asia (Desneux, Luna, Guillemaud & Urbaneja, 2011; EPPO, 2022; Mujica; Carhuapoma & Kroschel, 2022). In 2017, the pest was also confirmed to be present in the Xinjiang region of China and has become well established in northwestern and southwestern China (Zhang et al., 2021). *T. absoluta* is not only devastating tomato (*Lycopersicum esculentum* Mill.) production but also is becoming a more serious pest in potato (*Solanum tuberosum* L.) outside of its natural range of distribution with not only severe foliage damage and 50% yield reduction but also with direct tuber damage (Mohamed, Mohamed & Gamiel, 2012). The introduction of *T. absoluta* into Europe and its further spread is believed to be caused by the wide distribution and importance of its main host, tomato, which is traded within countries and among regions (Desneux et al., 2010). Further, the pest can develop on other widely cultivated Solanaceae such as potato, eggplant (*S. melongena* L.), sweet pepper (*Capsicum annuum* L.) and tobacco (*Nicotiana tabacum* L.) in addition to other non-cultivated Solanaceae (Smith et al., 2018; Mohamed, Mahmoud, Elhaj, Mohamed & Ekesi, 2015). This, in combination with other biological characteristics of the species such as the cryptic nature of larvae, high reproduction potential with multiple overlapping generations per year, and strong dispersal capacity likely supported its invasion of the old world (Urbaneja et al., 2013; Biondi, Guedes, Wan & Desneux, 2018). While its moderate to high resistance to commonly used insecticides (Guedes & Picanço, 2012; Guedes et al., 2019) is a growing concern, integrated pest management (IPM) programs are developed across different regions in which biological control via releasing or conserving arthropods natural enemies and sex pheromone-based control are the most promising and successful management practices (Desneux et al., 2022).

Numerous hymenopteran parasitoids in the families Braconidae, Eulophidae, Ichneumonidae, and Trichogrammatidae and predators in the families Miridae, Nabidae, Phytoseiidae among others are reported to attack *T. absoluta* in the new invaded areas of Europe, North Africa, and the Middle East (Zappalà et al., 2013), forming new pest-natural enemy associations with varying degrees of natural biological control. In Africa, the indigenous natural enemy diversity attacking *T. absoluta* is extremely scarce, and there is little evidence that new associations can substantially regulate the population of the pest. For example, in surveys carried out in different states of Sudan, no natural enemies of *T. absoluta* were detected (Mohamed et al., 2012). In Kenya, nine parasitoids were identified but with a combined parasitism rate of 7.26% only (Kinyanjui et al., 2021). In Tunisia, the mirid predator, *Nesidiocoris tenuis* (Reuter) (Hemiptera, Miridae) showed significant promise in suppressing populations of *T. absoluta* in open-field tomatoes when used in combination with pheromone lures, and selective insecticides (Indoxacarb and Spiromesifen) and bio-pesticides (*Bacillus thuringiensis* Berliner) (Abbes & Chermiti, 2010).

The absence of co-evolved efficient natural enemies in Africa might be the reason why *T. absoluta* population dynamics in the new invaded areas were explosive when compared with damages in the native area where natural enemies are diverse and common (Zappalà et al., 2013; Desneux et al., 2010). Desneux et al. (2011) emphasized that the knowledge on the distribution and natural biological control of *T. absoluta* in South America may permit pinpointing promising biocontrol agents for the pest. Hence, *T. absoluta*, being an invasive pest, and lacking efficient natural enemies in the new invaded regions, represents an excellent target for classical biological control. Challenges are to identify efficient biocontrol agents. The International Potato Center (CIP), Lima, Peru identified the koinobiont solitary larval endoparasitoid *Dolichogenidea gelechiidivoris* (Marsh) (Syn.: *Apanteles gelechiidivoris* Marsh) (Hymenoptera, Braconidae) as being highly effective in the natural range of *T. absoluta* distribution in Peru (Palacios & Cisneros, 1995). Upon request of the Kenyan Ministry of Agriculture, CIP compiled a dossier for the introduction of the parasitoid to Kenya according to the FAO Code of Conduct for the Import and Release of Exotic Biological Control Agents (FAO, 2017) and introduced the parasitoid in 2017 in collaboration with the International Centre of Insect Physiology and Ecology (*icipe*) to Kenya, after the approvals of the Kenya Plant Health Inspectorate Service (KEPHIS) and the Ministry of Agriculture of Peru to export the parasitoid to Kenya. Re-examining the diversity of *T. absoluta* parasitoids in its natural distribution of the Neotropics confirmed the importance of *D. gelechiidivoris* in Colombia, Chile, and Peru in controlling *T. absoluta* (Salas Gervassio, Aquino, Vallina, Biondi & Luna, 2019). As part of this first intended classical biocontrol program for *T. aboluta* in Africa, the potential of the parasitoid was further studied and confirmed (Aigbedion-Atalor et al., 2021; Mama Sambo, Ndlela, du Plessis, Obala & Mohamed, 2022; Ayelo et al., 2022), and recently released in East African countries (Ethiopia, Kenya, and Uganda) (Mohamed, Dubois, Azrag, Ndlela & Neuenschwander, 2022).

Temperature is one of the most important factors affecting the development, survival, and reproduction of insects and strongly determines demographic parameters, which are important for understanding population growth and dynamics, development rates and seasonal occurrence (Bale et al., 2002; Logan, Wollkind, Hoyt and Tanigoshi, 1976). For properly planning classical biocontrol programs, temperature-dependent pest, and natural enemy (parasitoid) phenology models are powerful analytical tools to be used in combination with Geographical Information Systems to map and analyze potentially suitable release areas and the efficacy of the parasitoid to control its host in the new environment (Kroschel et al., 2016; Cañedo, Dávila, Carhuapoma, Kroschel & Kreuze, 2022), which approach is applied in the Insect Life Cycle Modeling (ILCYM) software (www.cipotato.org/ilcym; Sporleder et al., 2013, 2020). Phenology models were developed for *T. absoluta* (Mohamed, Azrag, Obala & Ndlela, 2022; Mujica et al., 2022), and based on the phenology model developed for *D. gelechiidivoris,* Aigbedion-Atalor, Hill, Azrag, Zalucki & Mohamed (2022) concluded that inoculative augmentation can be considered in countries with temperature ranging between 15–30°C; however, a thorough analysis and mapping of suitable release areas globally has not been realized yet.

Hence, the objective of the present study was to develop and use an optimized temperature-dependent phenology model of *D. gelechiidivoris* for mapping and analyzing the potential establishment and efficacy of the parasitoid globally. We followed the approach we recently reported for the parasitoid *Apanteles subandinus* Blanchard (Hymenoptera, Braconidae), a larval endoparasitoid of the potato tuber moth, *Phthorimaea operculella* Zeller (Cañedo et al., 2022). Life table data of *D. gelechiidivoris* were collected at five constant temperatures in its main host *T. absoluta* to study the temperature-dependent development and survival of immature stages, complemented with data on the adult longevity and reproduction capacity derived from the publication of Aigbedion-Atalor et al. (2022). For model validation, simulated life table parameters were compared with observed life table data of the parasitoid collected at fluctuating natural temperature conditions. Finally, the *D. gelechiidivoris* phenology model was implemented in ILCYM’s potential population distribution and risk mapping module to map suitable release areas for the parasitoid using the Establishment Index (EI) and Generation Index (GI) displayed in those regions globally for which the potential establishment and distribution of its primary host *T. absoluta* is confirmed.

## Materials and Methods

### Origin and rearing of *T. absoluta*

A *T. absoluta* colony collected from different tomato production regions in Peru was maintained under laboratory conditions at the International Potato Center (CIP), Lima, Peru. One gram of tomato seeds was sown in plastic trays (50 x 30 x 30 cm) in a mixture of soil, sand, and moss (1:1:1 v/v/v) and placed in screenhouses under non-regulated temperature conditions. Seedlings were treated with fungicides (Acephate 4/1000, Vencetho Saume®) to prevent fungal diseases. After 7-10 days, individual tomato seedlings were transplanted into pots (10 x 10 cm) filled with Pro-mix® potting soil. At a plant stem height of about 30 cm, 15 pots were placed in wooden-framed cages (50 x 50 x 70 cm) covered with fine nylon gauze, in which 100 newly emerged *T. absoluta* adults of a mixed sex ratio were released for oviposition. A 5% sugar solution was provided as food on top of the nylon gauze. The oviposition cages were maintained at a temperature of 25°C ± 1°C,

>70% relative humidity, and a natural photoperiod of 12:12 (L:D) h. Three times, in a 3-day interval, the potted tomato plants were replaced, and the infested plants were stored in a separate rearing room at 25°C for 10-12 days, which allowed developing moths to complete their larval stage. Thereafter, plants were cut off at the bottom and laid in plastic containers with paper towel at the base. Pupae were collected after 7 days for continuing the rearing cycle or were used for the rearing or life table experiments of the parasitoid *D. gelechiidivoris*.

### Origin and rearing of *D. gelechiidivoris*

*D. gelechiidivoris* specimens used in this study were derived from a laboratory colony reared on *T. absoluta* on tomato plants maintained at the International Potato Center (CIP), Lima, Peru. Parasitoids were originally collected from *T. absoluta*-infested tomato leaves in the Cañete valley. Newly collected *D. gelechiidivoris* specimens from different coastal tomato production regions of Peru were made regularly to reduce any inbreeding effects (Haghani et al., 2006). In a 2-liter container, 20-30 newly emerged couples of parasitoids were released for mating for two days. A 5% sugar solution was provided as food on top of the nylon gauze. Tomato leaves infested with about 30-40 second instar of *T. absoluta* were conditioned in a small container (10 cc) with water to maintain plant turgor, in which fertilized females of *D. gelechiidivoris* were released. Infested leaves were replaced each 48 hrs, and leaves with parasitized larvae were transferred to a 2-liter plastic tray preconditioned with wet paper towel at the base. Non-infested tomato leaves were added as feed twice a week to ensure larval development of the pest. Emerged *D. gelechiidivoris* adults were extracted using an aspirator and used to restart parasitoid mass-rearing. Adults’ emergence occurred after 20-25 days at 22°C ± 2°C.

### Experimental procedure and data collection

The effect of temperature on the development of *D. gelechiidivoris* was studied in controlled incubator chambers (Thermo Fisher Scientific Inc., MA) at five constant temperatures of 10, 15, 20, 25, and 30°C. Data loggers (Hobo H8, Onset, MA) were used to monitor the temperature conditions. Relative humidity in the chambers was maintained at about 60% by placing containers with water; the photoperiod was kept at 12:12 (L:D) h.

#### Development of immature stages and survival

Four compound tomato leaves with at least 7 fully expanded leaflets were individually put in small glass tubes (5 cc) containing water and placed in a wooden cage (45 x 30 x 25 cm) covered with fine nylon mesh. Fifty 2-day-old *T. absoluta* adults of a mixed sex ratio were released in the cage for a 6-hr oviposition period. Egg-infested leaves were transferred to another wooden cage (45 x 30 x 25 cm) until the hatching of the eggs and the entrance of the first larval stage into the leaves. When the larvae reached the second larval instar, 30 couples of *D. gelechidiivoris*, after a mating period of 2 days, were released into the cage. After a 6 hr period of parasitism at the temperature used for general rearing (25°C), tomato leaves with at least 20 larvae were individually placed in Petri dishes and transferred to climatic chambers to study parasitoid development at the five constant temperatures.

At each constant temperature not less than one hundred *T. absoluta* larvae were inspected using the stereo microscope to determine *D. gelechidiivoris* parasitism rate and the development of eggs to larvae. To confirm parasitism and the development of eggs inside the larva, the dissection of *T. aboluta* larvae was performed as described by Cañedo et al. (2022). Larvae were observed until pupation to determine the larval development time as well as to record survival. *D. gelechidiivoris* pupae were maintained at the same constant temperature conditions until the emergence of adults. Adult emergence was recorded twice daily to determine development time of pupae and sex. This made it possible to evaluate total immature development time for both males and females of *D. gelechidiivoris*. In the life table, egg development time was included in the larva development time (egg-larva). The experiment was replicated three times at each constant temperature.

#### Adult longevity and reproduction capacity

In our life table experiments, it was not possible to conduct the studies on the adult longevity and reproduction capacity, thus estimates from bibliographic sources at constant temperatures were used from Aigbedion-Atalor et al. (2022). The data (means and standard errors) provided in this publication were ordered in a cohort-type life table and then subjected to simulations using R statistics (R Core Team, 2018). This procedure consisted of simulating the possible daily lifetime of adult’s females and males, and the oviposition capacity of females in a sample of 100 individuals. Additionally, the median time to oviposition was calculated based on the estimated time of effective oviposition during the entire lifetime of females. This variable was not considered by Aigbedion-Atalor et al. (2022).

### Data for model validation

The influence of fluctuating temperature on the development, mortality, and survival time of *D. gelechiidivoris* was studied under natural temperature conditions at the experimental station of CIP in La Molina, Lima (12° 05’ S, 76° 57 W, 250 m a.s.l.) from September to October 2017, following the same procedure as used in the constant temperature studies. A data logger was used to monitor the daily maximum and minimum temperature conditions.

### Model parameterization and analysis

The development of the *D. gelechiidivoris* phenology model and its life table parameter simulation was conducted using the Insect Life Cycle Modeling (ILCYM) software version 4.0 developed by CIP (Sporleder et al. 2020). The software is freely available at the institute website (https://ilcym.cipotato.org/) (Sporleder et al., 2017, Sporleder et al., 2013). Data obtained in the life table studies under constant temperature conditions were arranged in incomplete life table formats as required by the “model builder” of ILCYM to process, analyze, and develop the phenology model (development time and its variation, development rate, mortality, senescence, total oviposition, and relative oviposition frequency). The “validation and simulation” module of ILCYM was applied for simulating life table parameters and for model validation. The best-fit model was selected based on the Akaike Information Criterion (AIC), a well-known goodness-of-fit indicator (Akaike, 1973) or other built-in statistics (R^2^, Adjusted R^2^, MSE). The smaller the value of the AIC, the better the model fitted. For the selection of the best functions, statistical criteria and biological aspects of the species were considered.

#### Development time and its distribution

For development times and adult longevity, log-error distributions were assumed; the log-logistic, lognormal and Weibull model were tested as distribution link function, and the most appropriate distribution link function was chosen according to the maximum likelihood. ILCYM provides several different models that are adequate for describing the relationship between temperature and median development time, mortality, adult senescence, oviposition time and average fecundity per female. These functions generally are fitted in terms of rates (1/median time); however, in ILCYM and in this study, the functions were fitted in terms of ln-times. In addition, lower developmental thresholds and the thermal requirements for each life stage were calculated by means of linear regression between temperature and observed development rates (Campbell, Frazer, Gilbert, Gutierrez & MacKauer, 1974) using only data points within the linear range (data points at high temperature outside the linear range were deleted). The survival time of the immature stages of *D. gelechiidivoris* was calculated from the relative frequency of surviving test insects. Different nonlinear models (remodelled parabolic functions) available in the ILCYM package were adjusted by regression to describe the mortality rate in each life stage and fecundity by temperature. Development rate was expressed by the reciprocal of the mean development times for immature stages of *D. gelechiidivoris*. Mortality was calculated from the frequency of cohort mortality.

#### Simulation of life table parameters at constant temperatures

Life table parameters—i.e., net reproductive rate (R_0_), gross reproduction rate (GRR), intrinsic rate of increase (r_m_), finite rate of increase (λ), mean generation time (T), and doubling time (DT)—were estimated using the simulation tool in ILCYM (Southwood & Henderson, 2000; Sporleder et al., 2013). The estimates were based on the phenology model formed to simulate development, mortality, and reproduction of 100 individuals for a 1-yr period. Deterministic simulation was performed over a range of 10-30°C in 1° steps.

#### Model validation

The validation tool in ILCYM was applied to evaluate the ability of the developed phenology model to reproduce the *D. gelechiidivoris* life table data collected under fluctuating temperature conditions. Differences in development times, mortality rates and life table parameters namely, intrinsic rate of natural increase (r_m_), finite rate of increase (λ), and mean generation time (T), between simulated and observed life tables were statistically evaluated by using z-scores and t-statistics: z = (observed value−simulated value)/standard deviation of the simulated value. Experimental life table data were compared with model outputs produced by using the same temperature records as input data. The validation of the established model was done using deterministic simulation.

#### Mapping suitable release regions

For mapping suitable release regions, we implemented the presented *D. gelechiidivoris* phenology model in ILCYM’s potential population distribution and risk mapping module following the methodology described by Sporleder et al. (2017, 2020) and Kroschel et al. (2013), 2016). For analyzing the potential establishment and efficacy of *D. gelechiidivoris* to control *T. absoluta* globally, we used the Establishment Index (EI), Generation Index (GI), and Activity Index (AI) (Kroschel et al., 2016), and an index based on the differences in generations (ΔGI = GI^parasitoid^-GI^host^) developed annually by the parasitoid and its host *T. absoluta* as described by Cañedo et al. (2022). The four indices are simulated and displayed globally for those regions in which the potential establishment and distribution of *T. absoluta* has been confirmed (i.e., with an Establishment Risk Index [ERI] of >0.7, according to Mujica et al., 2022). For the spatial simulations we used temperature data for the year 2018 (CRU-TS 4.03) provided by Harris, Jones, Osborn and Lister (2014) and downscaled with WorldClim 2.1 (Fick & Hijmans, 2017) (https://www.worldclim.org/data/monthlywth.html).

### Complementary analysis

The linear degree-day model (thermal summation model; Campbell et al. 1974; Sporleder, Kroschel, Gutiérrez & Lagnaoui, 2004) was used to estimate the linear relationship between temperature and the rate of development of *D. gelechiidivoris*. The linear relationship is *Y* (=1/d) = *a* + *b*T, where, *Y* is the rate of development (1/d), *T* is the ambient temperature (°C), and the regression parameters are the intercept (*a*) and slope (*b*). The thermal constant K (= 1/b) is the number of degree-days above the threshold summed over the development period. The theoretical lower (minimum) development threshold T_min_ (= -*a*/*b*) is the minimum temperature at which the rate of development is zero or no measurable development occurs. Data on development time of the different life stages, adult longevity, and fecundity of females were compared across constant temperatures using a one-way ANOVA of the statistical package R 4.1.0. When significant differences were detected, multiple comparisons of treatments were made using the Tukey test (P ≥ 0.5).

## Results

### Development and its distribution

The duration of the immature stages and the time required for *D. gelechidiivoris* to complete its life cycle from egg to adult decreased significantly with increasing temperatures within the temperature range from 15 to 30°C (Table 1). The parasitoid successfully completed its development from egg to adult in all temperatures evaluated, except at 10°C, which was lethal to pupae. Total development was almost four times longer at 15°C (52.8 d) than at 30°C (14.2 d) respectively. The log-logistic distribution model produced the best fit, based on the lowest AIC values, to describe variability in the development from egg-larvae and pupae according to temperature (Table 1, Figure 1). The common slopes determined for each life stage were highly significant (P < 0.001) and adequate to describe the overall variability in the development within each immature life stage (Table 1).

**Fig. 1.**
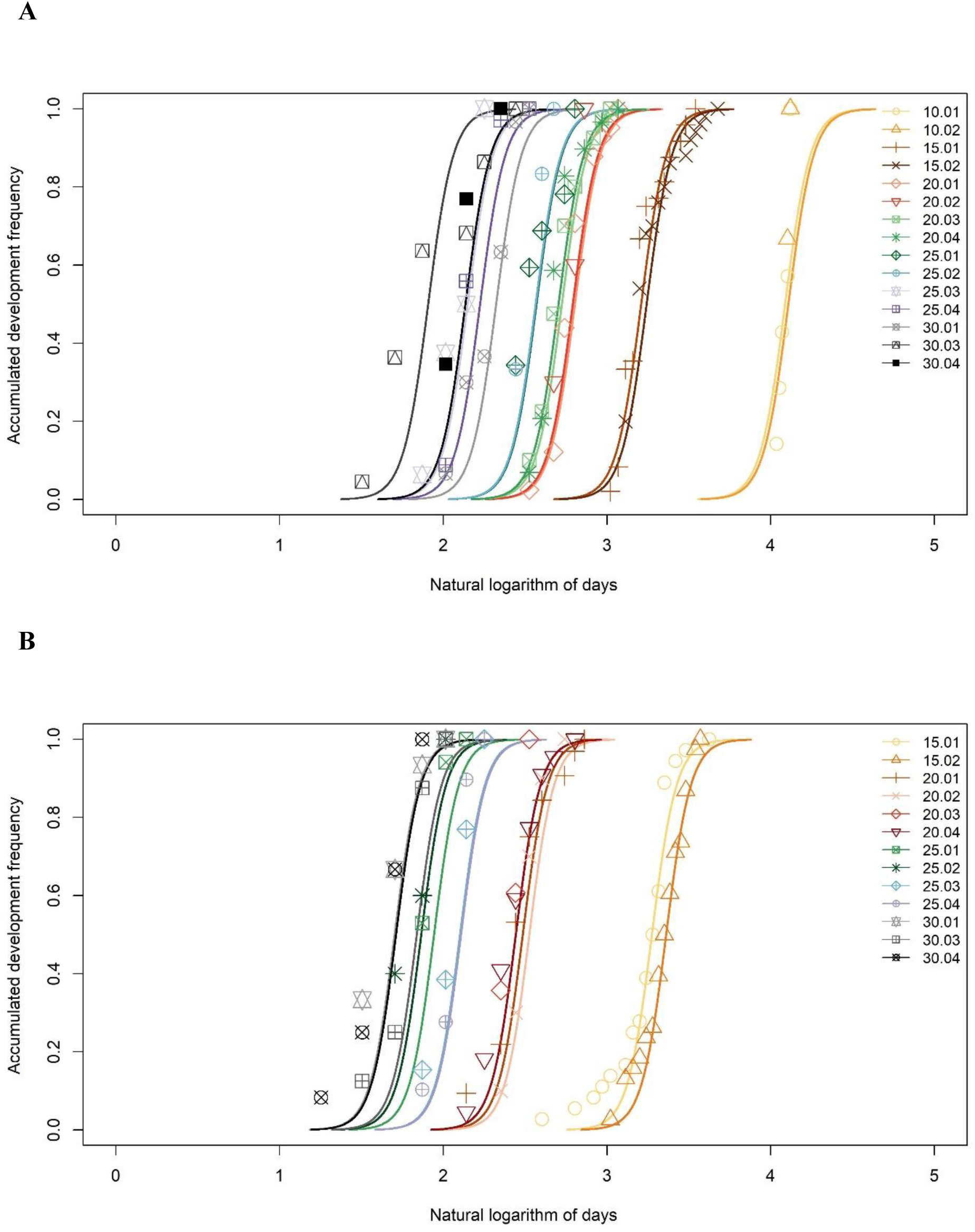
Cumulative distribution of development times of immature *D. gelechiidivoris* life stages (A: egg-larvae, B: pupae), reared on *Tuta absoluta* at constant temperatures. Fitted curves: log-logistic model.

### Development rate

The Janish-1 model described satisfactorily the temperature-dependent median developmental rate for the egg-larvae and pupae immature stages (Table 2). The coefficient of determination (R^2^) values explained 94% and 97% of the variation in median development times for the egg-larvae and pupae immature stages, respectively. The curves for the immature stages were linear from 15°C to 25°C, and consequently, a linear model would be satisfactory to describe temperature-dependent development between these temperatures (Figure 2). However, above 25°C development is curvilinear such that the Janish-1 model was more appropriate, as corroborated by the high R^2^. Estimated theoretical lower threshold temperature (estimated from the slope and intercepts of the linear regression) for the development of immature stages were 7.6°C (egg-larvae), 10.9°C (pupae), and 9.6°C (total immature). According to these thresholds, the thermal constant (k) for the development, expressed in degree-days (DD = 1/ slope), was 192.3, 108.7, and 294.1 for egg-larvae, pupae, and total immature stages respectively.

**Fig. 2.**
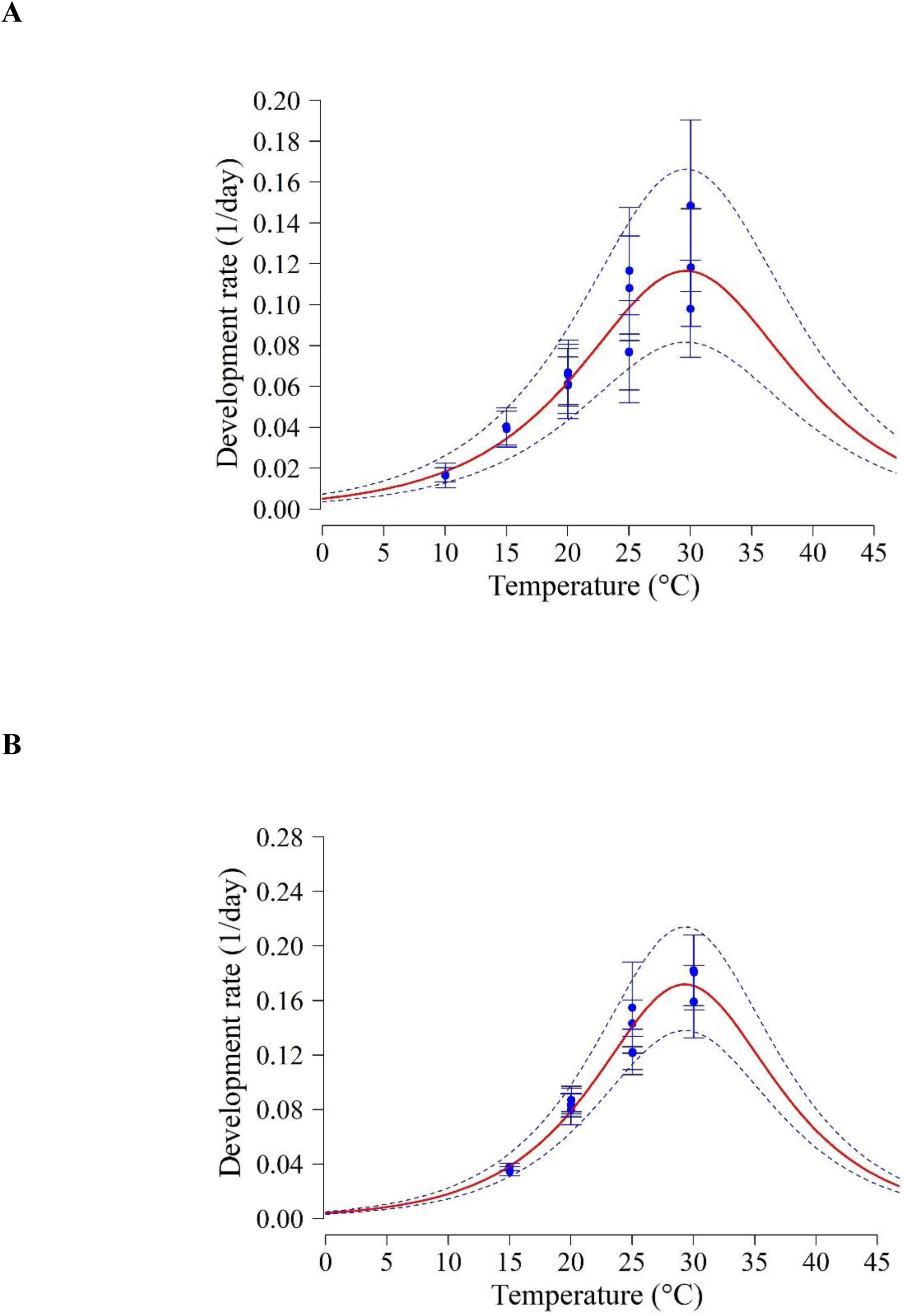
Temperature-dependent development rates for immature life stages of *D. gelechiidivoris*. The Janish-1 model for egg-larvae (A) and pupae (B) (solid line). Dashed lines represent the upper and lower 95% confidence limits. Bars represent standard deviation of the mean development rate.

### Immature mortality

The effects of temperature on the mortality rate of *D. gelechidiivoris* immature stages were best described by the Quadratic model for the egg-larvae and pupae stages (Table 3, Figure 3). The total immature mortality was lowest at 20°C (26%) and highest at 30°C (80.5%) (Table 1). At 10°C, all pupae died (before adults could emerge). The egg-larval and pupae stages had the lowest mortality at 15°C (18.4%) and 20°C (15.4%), respectively. The models explained >69% of the variation by temperature. Nonlinear models predicted optimal temperature for immature survival between 20– 25°C (26–33% mortality of all immature stages).

**Fig. 3.**
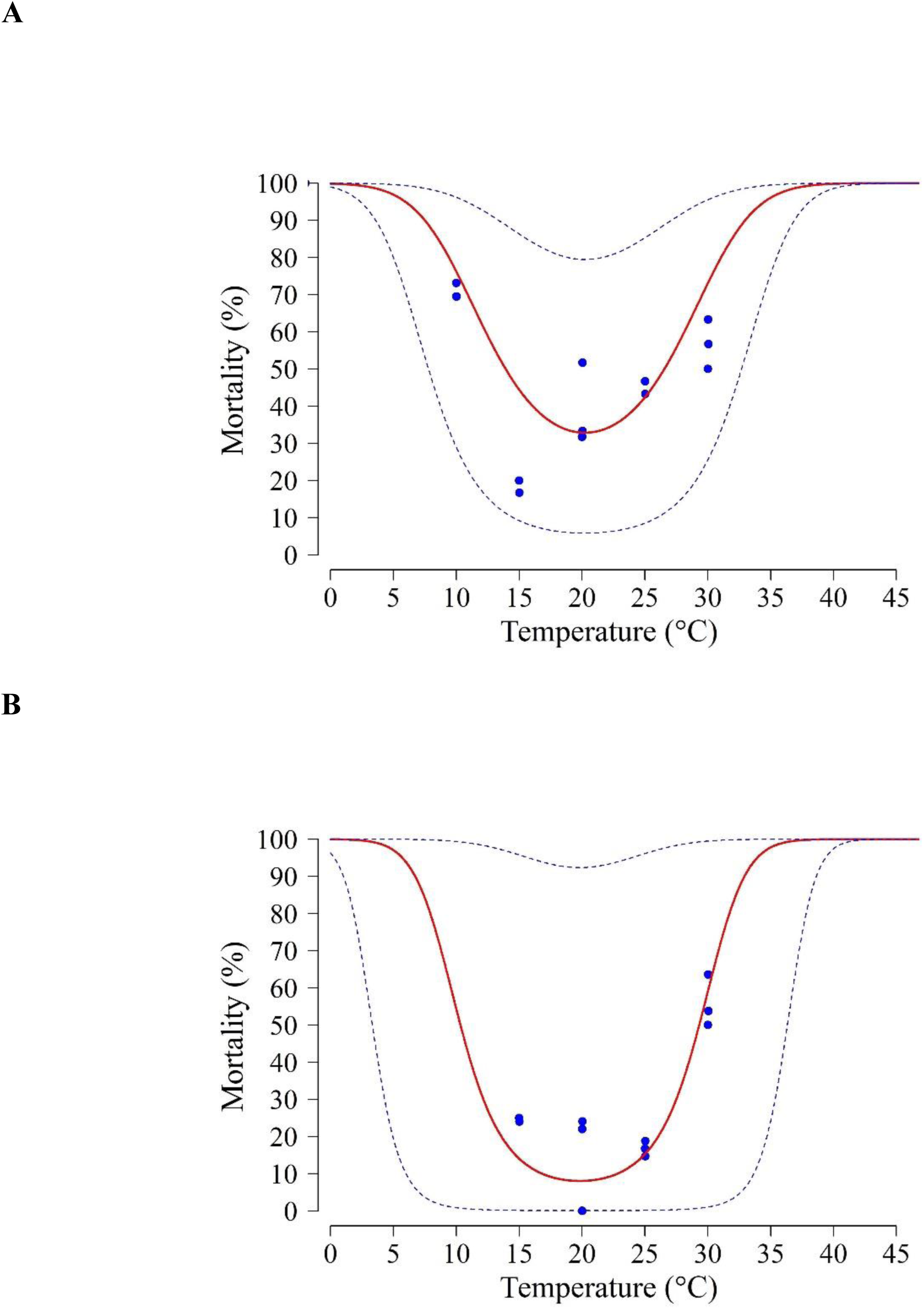
Temperature-dependent mortality rates of immature life stages of *D. gelechiidivoris*: (A) egg-larvae, (B) pupae. Fitted curves: Quadratic model.

### Adult longevity and fecundity

The longevity of adult female and male *D. gelechidiivoris* decreased with increasing temperature (Table 4), with significant differences between temperatures. Longevity of females ranged from 36.4 (10°C) to 3.6 (35°C) days and of males from 29.3 (15°C) to 3.6 (35°C) days respectively. A Quadratic model was fitted to determine the relationship between senescence rate of female and male adults and temperature (Table 5). The lowest senescence rates for females and males were observed within the temperature range of 10–20°C (Figure 4).

**Fig. 4.**
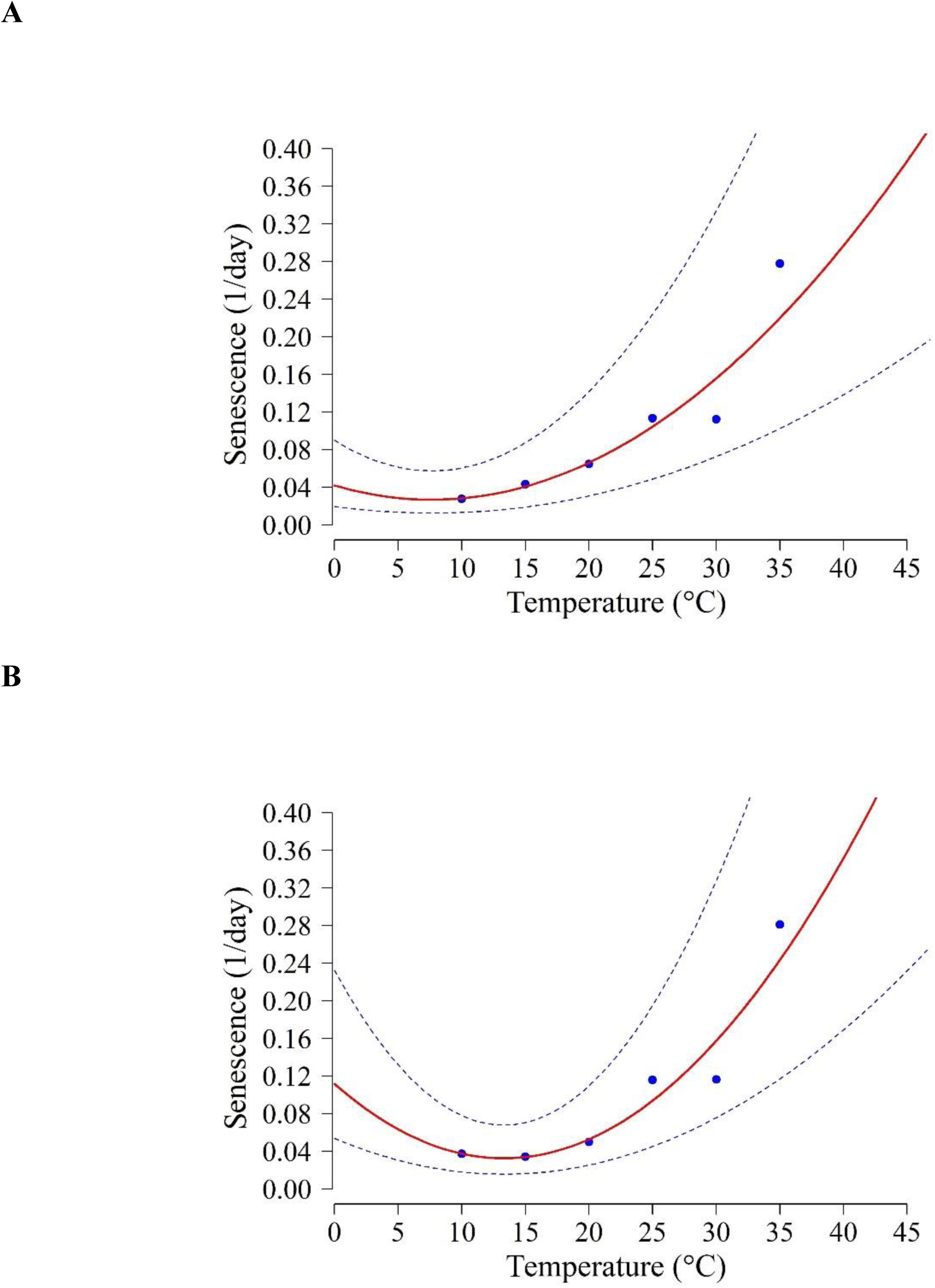
Temperature-dependent senescence rates for *D. gelechiidivoris* adult females (A) and males (B). Fitted curves: Quadratic model (solid lines). Dashed lines represent the upper and lower 95% confidence limits. Bars represent standard deviation of the mean senescence rate.

Median oviposition time decreased significantly with increasing temperature from 16.7 days at 10°C to 1.62 days at 35°C (Table 4). The relationship between temperature and oviposition rate was best described by a quadratic model (Table 5, Figure 5b). The effects of temperature on fecundity were best described by the Taylor model with predicted highest fecundity at 20°C (Table 4, Figure 5a). Fecundity per female was variable, ranging from 12.5 eggs at 35°C to 74.4 eggs at 20°C (Table 4). A Tukey test revealed significant differences between fecundity across all temperatures (F = 9591.94; df = 5,22531, p < 0.0001). The relationship between the cumulative proportion of progeny produced per female and normalized female age was described by the Weibull function (Table 5, Figure 5c). At 20°C, 50% oviposition was completed by the time the female reached a normalized age of 1.8.

**Fig. 5.**
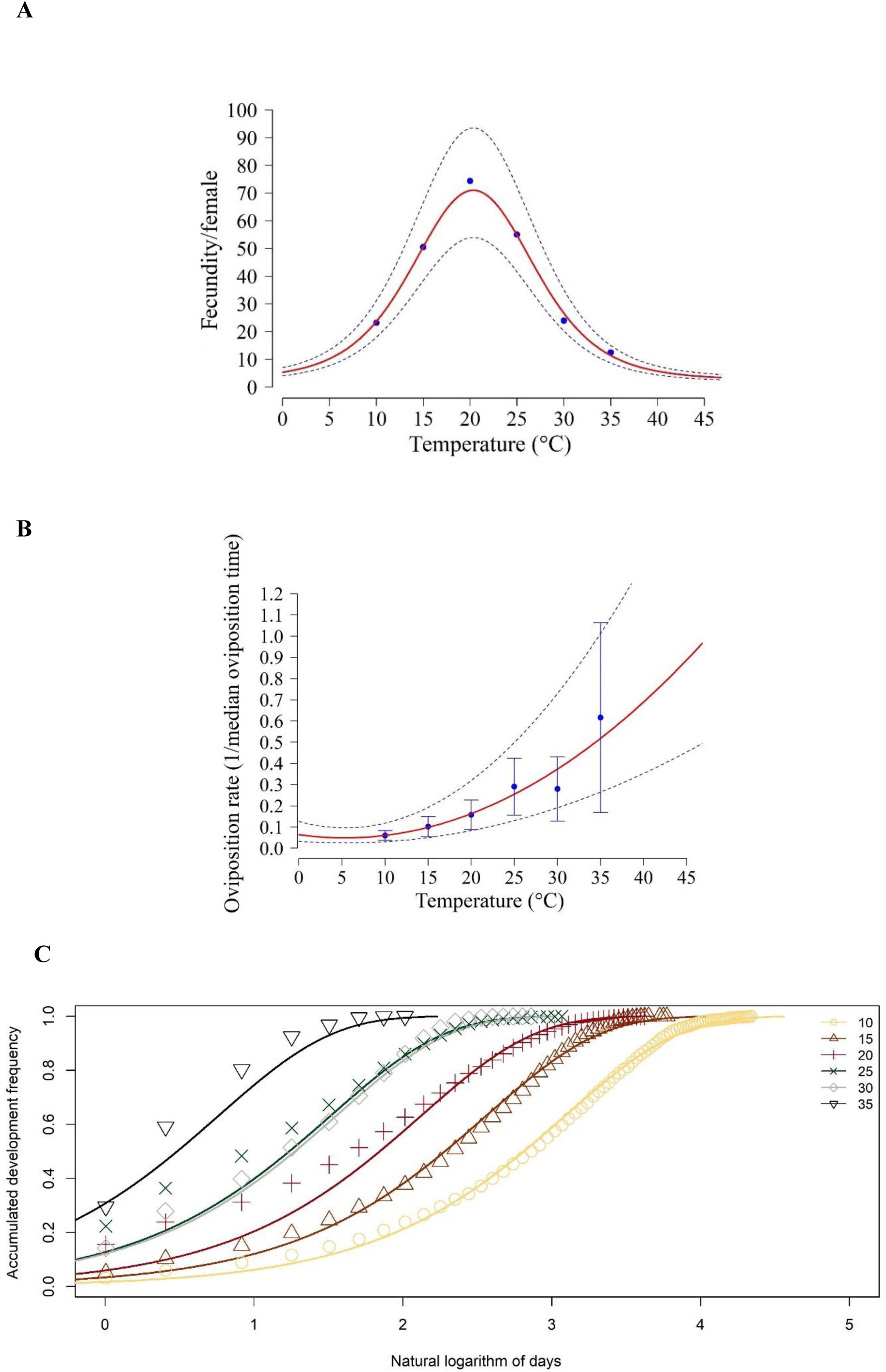
Fecundity of *D. gelechiidivoris* and its dependence on temperature and female age. (A) total fecundity per female, (B) temperature-dependent oviposition rate and (C) accumulated reproduction frequency. In (A) and (B) dots are observed data, solid lines are fitted models and broken lines are 95% confidence limits for the fitted model. In (b) bars represent standard deviation of the oviposition rate.

### Life table parameters

Simulations of *D. gelechidiivoris* population parameters showed that the intrinsic rate of natural increase (r_m_) augmented almost linearly with increasing temperature representing an asymmetrical dome-shaped pattern to reach a maximum at 24.4°C (0.1033) and decreasing sharply at 30°C (0.0276), with a minimum value at 12°C (0.0119) (Figure 6A). Similarly, the finite rate of increase peaked at 25°C (λ of 1.1077) and was smallest when exposed to 12°C (λ of 1.012) and 29°C (λ of 1.028; Figure 6B). The highest values for the gross reproductive rate (GRR) (Figure 6C) and net reproductive rate (R_0_) (Figure 6F) were found at 20°C with 49 offsprings/female and 19 females/female, respectively. The shortest mean generation time (T) was observed at 30°C (19.9 days) (Figure 6D) and the shortest doubling time (Dt) was determined at 24°C (6.84 days) (Figure 6E).

**Fig. 6.**
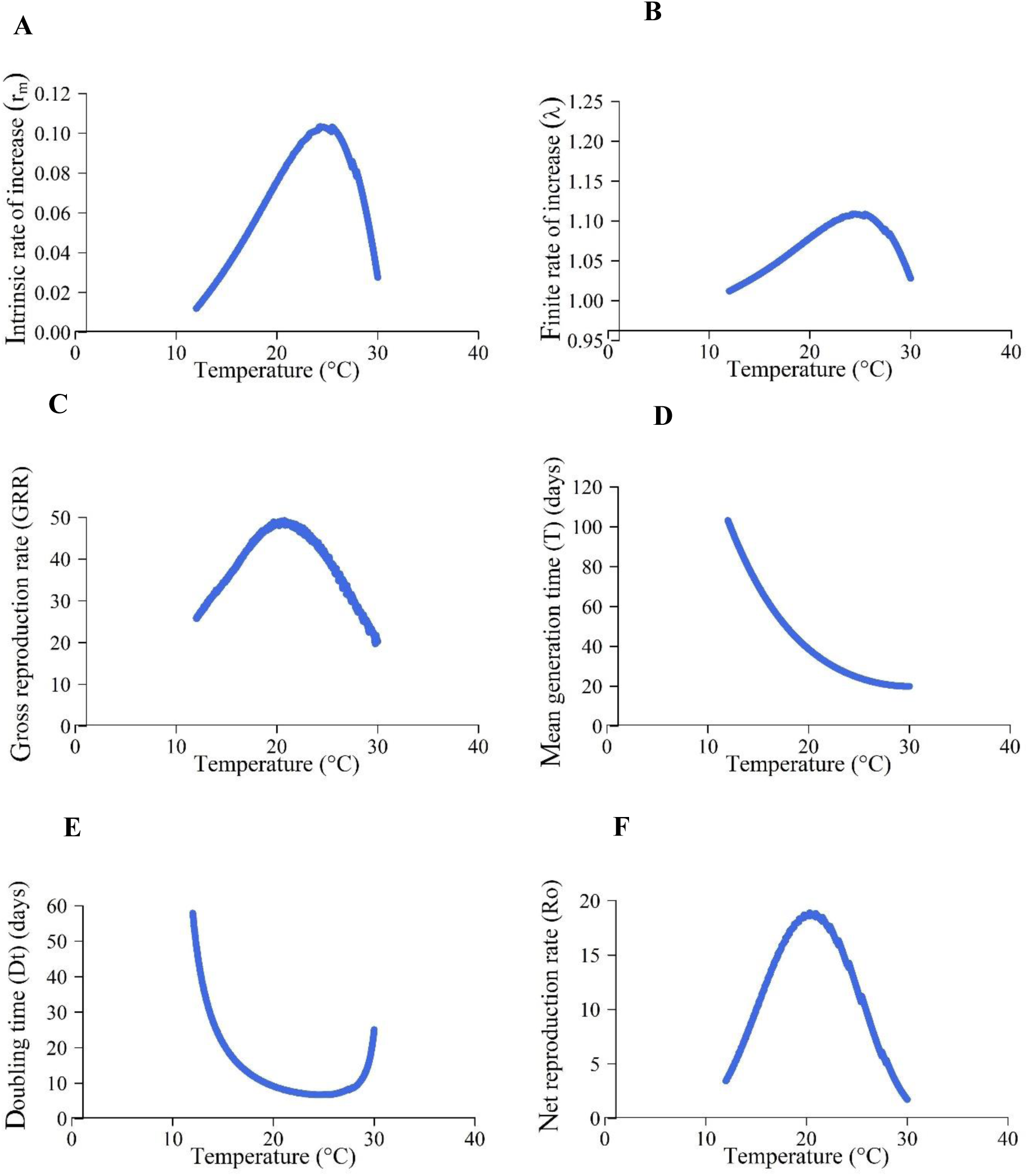
Simulated life table parameters of *D. gelechiidivoris* using the phenology model at constant temperatures. (A) Intrinsic rate of natural increase (r_m_), (B) finite rate of increase (λ), (C) gross reproduction rate (GRR), (D) mean generation time (T), (E) doubling time (Dt) and (F) net reproduction rate (R_o_).

### Validation of the model

Phenology model validation of *D. gelechidiivoris* was carried out using fluctuating temperature data at a range from 12-29°C, with an average mean temperature of 17°C. Simulated population parameters like intrinsic rate of natural increase (rm), finite rate of increase (λ), and mean generation time (*T*) were well predicted when compared with observed data collected under fluctuating temperature (Table 6). Simulation predicted 15% lower development time from egg to adult (28.8 d) compared with data collected at fluctuating temperatures (34.1 d).

### Suitable release regions

An Establishment Index (EI)=1 indicates survival and reproduction of *D. gelechidiivoris* throughout each day of the year, which means that the likelihood of long-term establishment for classical biological control is very high (Figure 7). Regions with a very high potential for successful releases (EI>0.9) in countries invaded by *T. absoluta* are in Central and South America, mostly all parts of Africa, Southern Asian countries, the Middle East Asia, and Southern European countries. This high establishment potential is associated with a Generation Index (GI)>13 (up to 20 generations) in tropical and >6–13 generations in subtropical regions, which are potentially developed by *D. gelechidiivoris* per year (Figure 8). Moreover, the number of generations developed by *D. gelechidiivoris* per year surpass the generation numbers of its host *T. absoluta* by >2.4–3.3 generations in tropical and >1.6–2.4 generations in subtropical regions, respectively, indicating an overall good biocontrol potential and capacity of *D. gelechidiivoris* in these regions (Figure 9). The GI is strongly correlated with the activity index (AI) (Figure 10). The AI indicates most appropriately the potential population growth averaged throughout a year; each increase of the AI by 1 indicates a 10-fold higher increase rate of the population. An AI of 6–13 is predicted for subtropical regions, and of <13–21 for tropical regions. In regions where temperatures have reached the upper temperature threshold, i.e., >30°C for the development of *D. gelechidiivoris,* the generation number will still increase, but the population growth will gradually reduce due to increasingly high temperature-induced mortality and reduced reproduction per female (Table 1, Table 4).

**Fig. 7.**
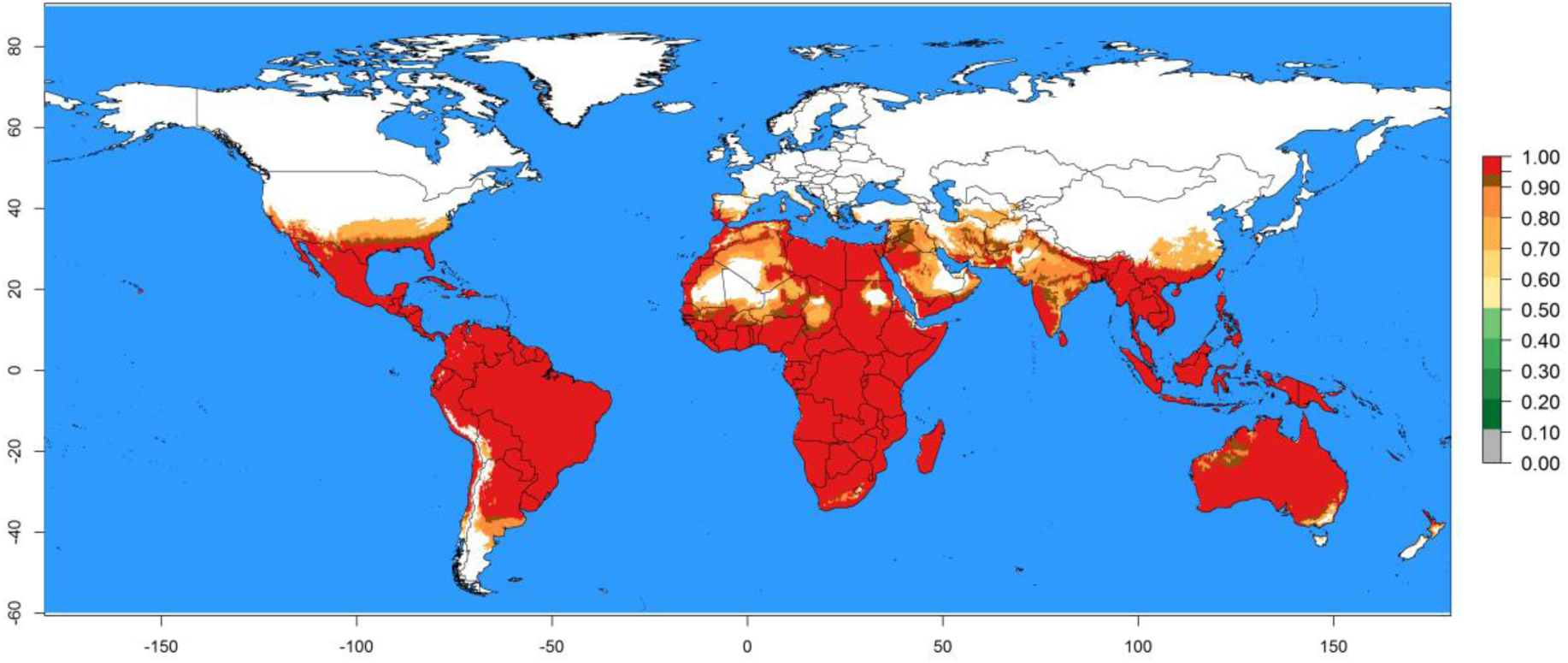
Potential establishment and distribution of *D. gelechiidivoris* according to model predictions, using the Establishment Index (EI) for the year 2018 globally for which the potential establishment and distribution of its primary host *T. absoluta* is confirmed (i.e., with an Establishment Risk Index (ERI) >0.7).

**Fig. 8.**
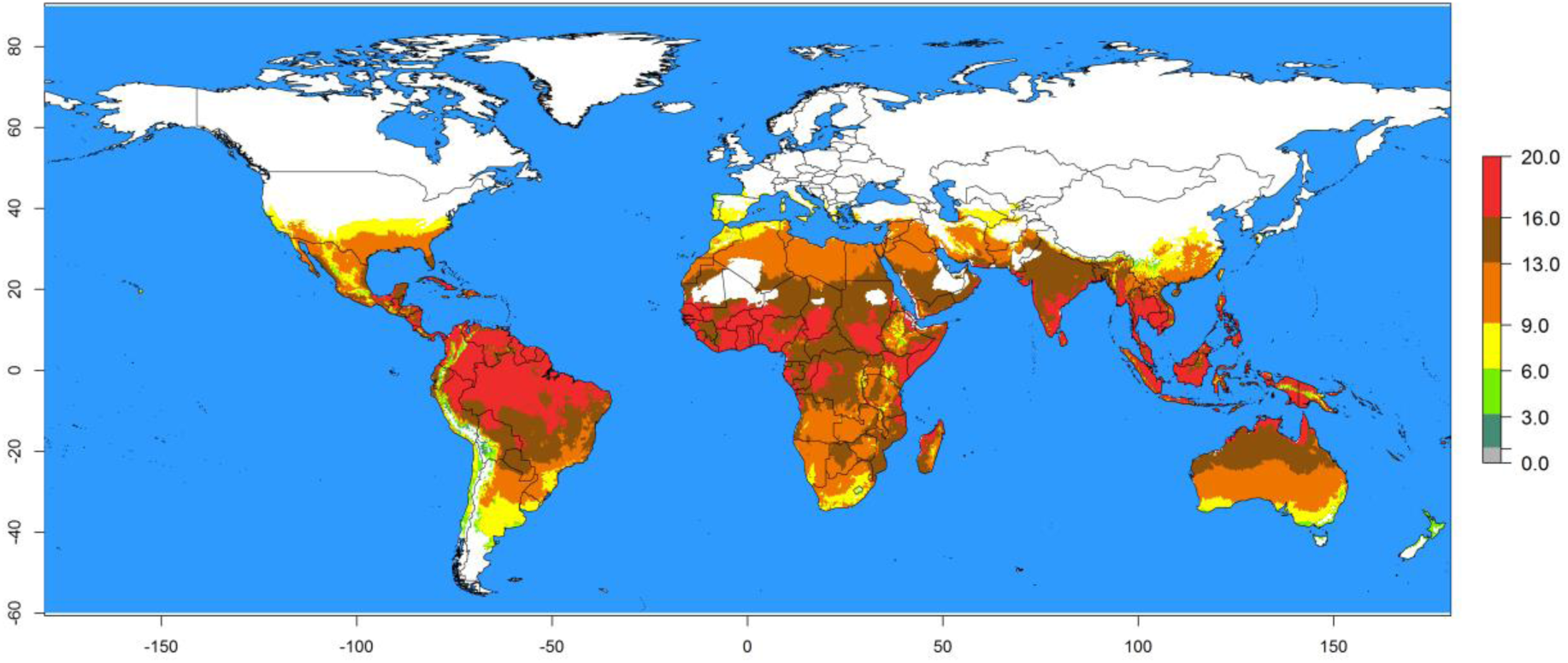
Potential abundance of *D. gelechiidivoris* according to model predictions, using the Generation Index (GI, number of generations/year) for the year 2018 and displayed globally for which the potential establishment and distribution of its primary host *T. absoluta* is confirmed.

**Fig. 9.**
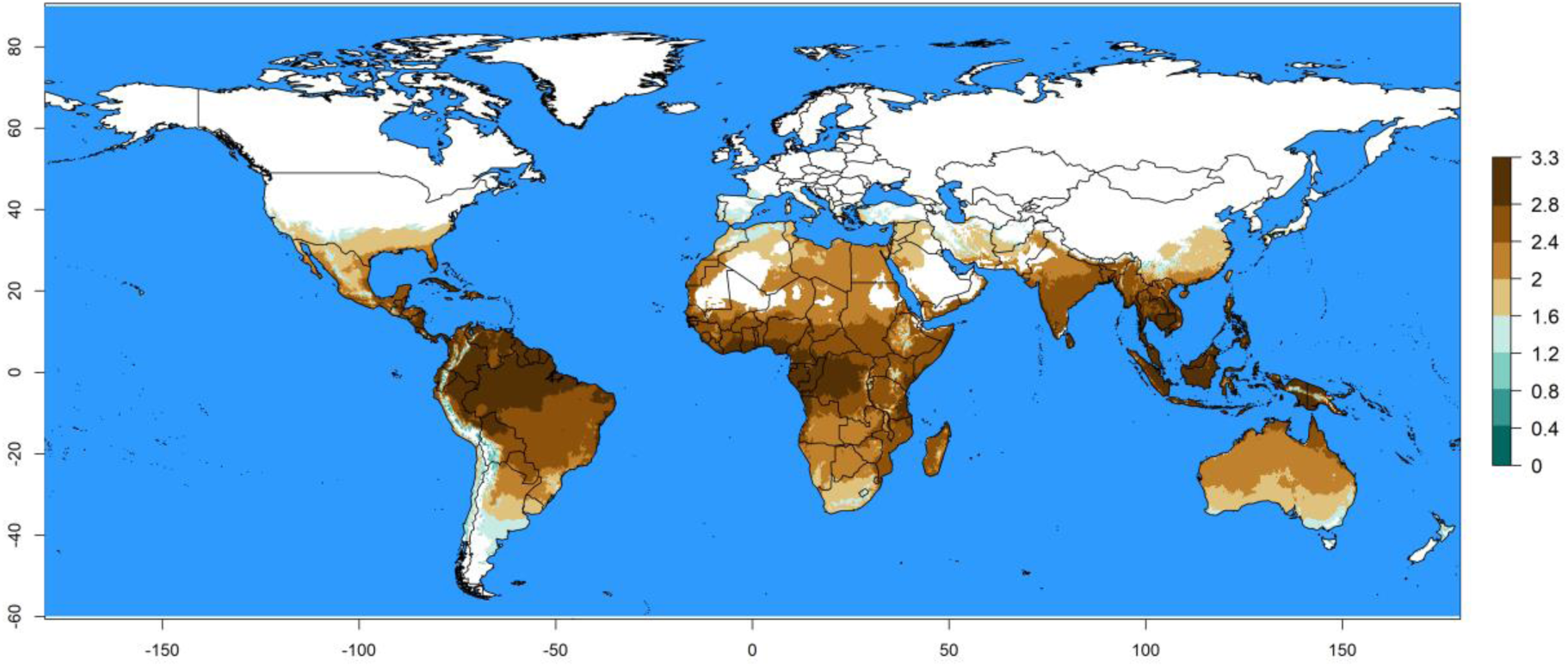
Potential efficacy of *D. gelechiidivoris* to control *T. absoluta* according to model predictions using the differences in the annual number of generations developed by the parasitoid and its primary host (ΔGI= GI^parasitoid^-GI^host^) for the year 2018 and displayed globally for which the potential establishment and distribution of *T. absoluta* is confirmed. Higher ΔGI values indicate larger biocontrol capacity.

**Fig. 10.**
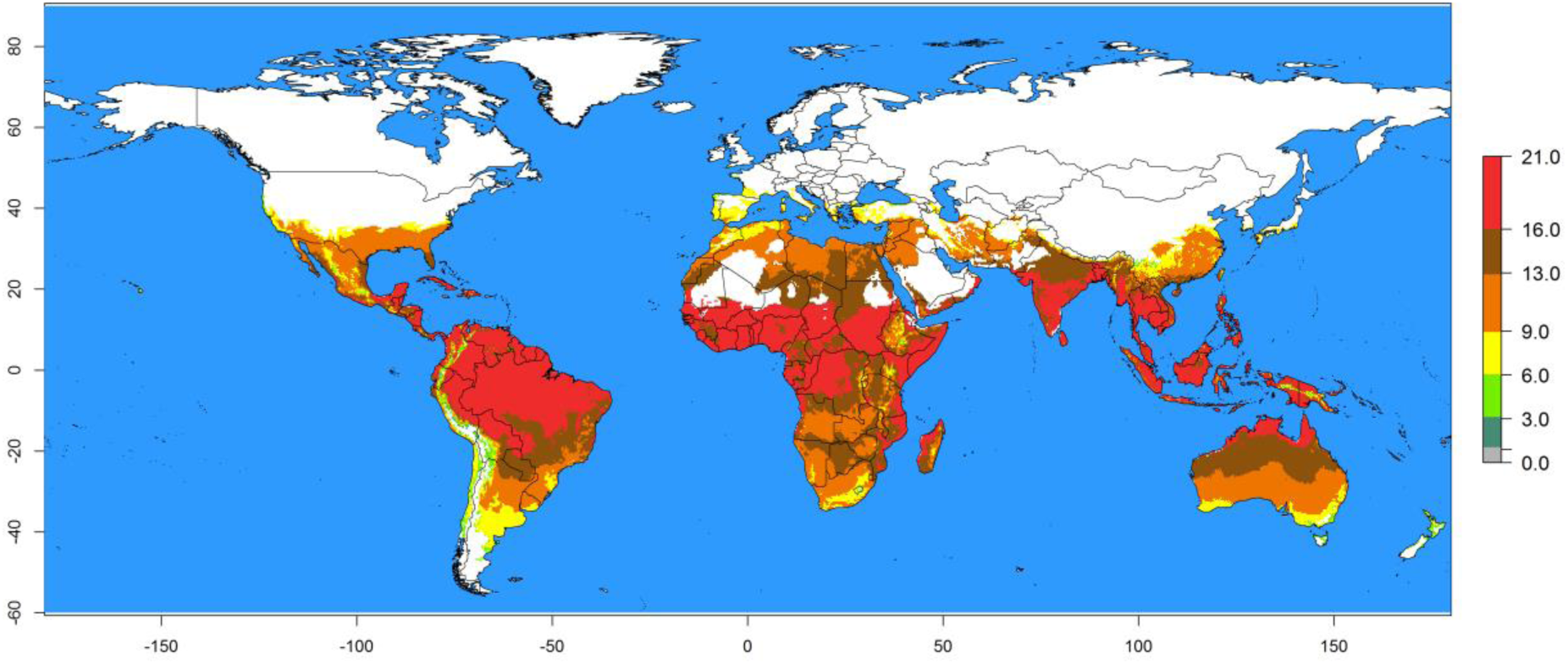
Activity of *D. gelechiidivoris* according to model predictions, using the activity index (AI, potential population growth) for the year 2018 and displayed globally for which the potential establishment and distribution of its primary host *T. absoluta* is confirmed. An activity index of 1 indicates a 10-fold higher increase rate; i.e., an index of 4 indicates a 10,000-fold (=10^4^) potential increase of population numbers within one year.

## Discussion

The koinobiont oligophagous larval endoparasite *D. gelechiidivoris* has a wide ecological amplitude in its native range of distribution in the Neotropics where it is adapted and occurs at coastal regions of Peru, Colombia, and Chile as well as in the Andean Highlands (Palacios & Cisneros, 1995; Vallejo, 1999; Larrain, 1987) as parasitoid of *T. absoluta,* and the potato tuber moths, *Phthorimaea operculella* and *Symmetrischema tangolias* (Gyen). Although in potato agroecosystems of the Peruvian Andean Highlands only few individuals have been recovered parasitizing *P. operculella* at altitudes between 2800 and 3850 m asl, it demonstrates that the species is adapted to these harsh temperature conditions with only one potato growing period from November to May (Kroschel & Cañedo, 2009). In the central Peruvian lowlands at the coast of Peru in the Cañete valley, *D. gelechiidivoris* represented 14% of the total parasitoid guild collected in untreated potato fields (Mujica & Kroschel, 2013). Periodic field sampling at the central Coast of Peru showed that *D. gelechiidivoris* was the predominant endoparasitoid of the potato tuber moth with 26-41% of parasitism in farmers’ fields (with application of insecticides) and 57% of parasitism in an isolated field without the use of insecticides (Palacios & Cisneros, 1995; Cañedo & Cisneros, 1994). It has also been reported as the predominant parasitoid of *T. absoluta* in La Molina, Lima and the Rimac valley (Redolfi de Huiza & Vargas, 1981; Sanchez & Redolfi de Huiza, 1985). In Colombia, *D. gelechiidivoris* showed a high potential as biological control agent to effectively reduce *T. absoluta* populations in tomato with parasitism rates of 77% (Vallejo, 1999). Accordingly, biological control in tomato allowed keeping *T. absoluta* as a secondary pest at non-economic damage levels, thereby increasing crop productivity while reducing the use of chemicals (García, 2000).

The biocontrol potential of *D. gelechiidivoris* has been already previously used in classical biocontrol of *T. absoluta* in Chile (Rojas, 1997), of *P. opercullela* and the tomato pinworm, *Keiferia lycopersicella* (Walsingham) in California (Marsh, 1975) as well as of *K. lycopersicella* in Hawaii (Nakao & Funasaki, 1997). In Chile, *D. gelechiidivoris* established in the central region (Estay & Bruna, 2002) but high levels of parasitism were first recorded after 10 years of the initial introduction (Desneux et al., 2010). The introduction and release of *D. gelechiidivoris* on the Eastern Island was successful and highly effective reducing the *T. absoluta* infestation (Ripa & Rojas, 1984; Ripa, Rojas & Velasco et al., 1995).

Several studies were conducted to better understand and rate the biocontrol efficacy of *D. gelechiidivoris* based on various biological parameters. Van Lenteren et al. (2021) ranked the biocontrol efficacy of *D. gelechiidivoris* as low in tomato production with *T. absoluta* as the sole pest species without the provision of additional food as the population development is considered slow with a limited capacity to parasitize and kill its host, which is in contradiction to the good biocontrol efficacy of the parasitoid reported from countries in South America. Laboratory studies on the performance of *D. gelechiidivoris* demonstrated 55% parasitism of early larval instars of *T. absoluta,* which was suggested to confirm its efficacy for biocontrol programs (Aigbedon-Atalor et al., 2020). The study of Mama Sambo et al. (2022) showed a functional response type II by *D. gelechiidivoris,* which suggests density-dependent parasitism up to a certain level of host density at which the attack remains constant regardless of the increase in host density (Holling, 1959). Type II functional response is suggested to be satisfactory for regulations of pest populations whereas type III is classified as ideal (Fernández-arhex & Corley, 2003). In solitary larva parasitoids, such as *D. gelechiidivoris*, only one larva per host completes its development. Many solitary parasitoids have therefore evolved mechanisms to avoid superparasitism as this would result in intraspecific competition between progenies and reduce the profits of oviposition. Mama Sambo et al. (2022) showed that superparasitism was very low by *D. gelechiidivoris* and generally occurred at the lowest host densities which was suggested to be another good indicator for its efficacy. The same authors also found a high emergence rate of wasps of 34% and 51% for single and group foraging females respectively. In comparison, an emergence rate of 53% was reported at a higher parasitoid-host ratio of 1:20 by Aigbedon-Atalor et al. (2020). Although Mama Sambo et al. (2022) found a slightly male-biased sex ratio when *D. gelechiidivoris* was exposed in a group to *T. absoluta* larvae, this was not confirmed in the study of Aigbedon-Atalor et al. (2020). Also, Bajonero et al. (2008) reported a higher female bias at temperatures between 14-26°C, being highest at higher temperatures. The only study which was conducted to understand the efficacy of releases of *D. gelechiidivoris* was reported by Wanumen (2012) in Colombia, who estimated a ratio of 6:1 (third instar larvae of the pest:parasitoid female) for successfully implementing biological control of *T. absoluta* under greenhouse conditions.

The determination of the parasitoid’s temperature-dependent development is crucial for better predicting the potential of the parasitoid to establish in new regions and to control the target pest. In a preliminary study, Bajonero et al. (2008) demonstrated that *D. gelechiidivoris* can complete its life cycle at temperature ranging from 14-32°C. More recently, Aigbedon-Atalor et al. (2022) studied the parasitoid development at constant temperatures between 10-35°C and reported the completion of the life cycle between 10-30°C. At 35°C, no development from egg to larva was found. In our study, *D. gelechiidivoris* completed the life cycle at constant temperatures from 15-30°C but 10°C was lethal to pupae. Although Aigbedon-Atalor et al. (2022) confirmed 10°C as the lower threshold for development, a high mortality of >90% occurred at this temperature. The established functions describing the temperature-dependent development, survival, and oviposition allowed the development of an overall *D. gelechiidivoris* phenology model. Life table parameters simulated at constant temperatures indicated that *D. gelechiidivoris* population develops within the temperature range of 12-30°C, with an optimum temperature between 20-25°C (with the lowest mortality of 26–33% of all immature stages), which are consistent with the findings by Bajonero et al. (2008) and Aigbedon-Atalor et al. (2022). The intrinsic rate of increase, which indicates the most favorable temperature for population growth (i.e., development time, survival, and reproduction; Southwood & Henderson, 2000) reached its highest value at 24.4°C. Also, at 24°C, the finite rate of population growth was highest and doubling time shortest. Finite rate of population growth and doubling time are the most important parameters describing population increase. In our study, the theoretical lower threshold temperature for the development of egg-larvae, pupae, and total immature stages of *D. gelechiidivoris* were 7.6°C, 10.9°C, and 9.6°C, respectively. These results are like those found by Aigbedon-Atalor et al. (2022) for the egg-larvae stage (7.02°C) but differ for the pupal stage (5.93°C). Lower threshold temperatures for the egg-larvae stage can be explained to the fact that these stages develop inside its host. *T. absoluta* larvae act as thermal barrier for the first developmental stages of the parasitoid, while pupae are fully exposed to temperature of the environment (Bajonero et al., 2008).

Our mapping of suitable release regions confirms the high adaptability of *D. gelechiidivoris* to establish in a wide range of environments in tropical and subtropical regions, which have been successfully invaded by *T. absoluta* causing significant economic damage in tomato fields. An Establishment Index (EI) of 1 indicates regions with the highest establishment potential of the parasitoid and a population growth throughout the year (Kroschel et al., 2013, 2016). An EI of 1 has been confirmed for the parasitoid’s native range of distribution and where it was released by former biocontrol programs, e.g., in Chile or California. Recently, the occurrence of *D. gelechiidivoris* was reported for the first time in Europe in commercial tomato crops in Catalonia (North-eastern Spain) with the first detection of the parasitoid in larvae samples of the year 2016 (Denis et al., 2021). Although in the first years of detection only few individuals were found, parasitism rates steadily increased from May (2.7%) to October (21.8%) in 2020, indicating that the parasitoid has well established in the region. Likewise, the detection and occurrence of *D. gelechiidivoris* has been reported from tomato fields in several regions of Algeria with sampling starting in 2020 (Krache et al., 2021). The introduction into Spain and likely from Spain into Algeria occurred probably unintentionally from the Neotropics already some years ago because of global trade, which is also the assumed cause of arrival of *T. absoluta* in Spain (Desneux et al., 2010). Interestingly, in north-eastern Spain, the establishment of *D. gelechiidivoris* has been now confirmed to occur in a region for which an EI>0.7 has been determined, which indicates the potential of the parasitoid to survive in a region in which the population growth is restricted to certain periods of the year (like in the Andean region of Peru as reported by Kroschel & Cañedo, 2009), and with this value during a 9-months period only. In comparison, the establishment in Algeria along the Mediterranean coast occurred in regions with an EI=1. For the releases of *D. gelechiidivoris* in the East African countries of Ethiopia, Kenya, and Uganda, recently implemented by the International Centre of Insect Physiology and Ecology (*icipe*) as part of a classical biocontrol program for *T. absoluta,* the establishment of the parasitoid could be already confirmed (Mohamed et al., 2022); the high potential for establishment with an EI=1 was predicted for all three countries. Cañedo et al. (2022) introduced an indicator for the potential control efficacy which is expressed as the difference of generations (ΔGI) of the parasitoid and its host, which are developed per year. According to the developed maps, potentially suitable release areas for *D. gelechiidivoris* with a high control efficacy are tropical regions but releases of *D. gelechiidivoris* can be also considered for subtropical regions. This is demonstrated by a similar control efficacy determined for subtropical regions of South America, in which the parasitoid had been successfully released and used in biocontrol programs, and countries of southern Europe and North Africa.

In conclusion, the developed phenology model and simulated life table parameters estimated for *D. gelechiidivoris* reflect the temperature-dependent growth potential of the parasitoid in its main host *T .absoluta*. The phenology model of the parasitoid was successfully applied in the GIS-based application of the ILCYM software to generate global maps based on four indices (EI, GI, ΔGI, and AI) to predict suitable release areas based on temperature (not considering other abiotic factors such as precipitation or humidity). This approach should support and increase the successes of classical biological control programs using the koinobiont oligophagous larval endoparasite *D. gelechiidivoris* to control *T. absoluta*.

## Acknowledgements

This research was undertaken as part of, and partially funded by, the CGIAR Research Program on Roots, Tubers, and Bananas (RTB) and supported by CGIAR Trust Fund contributors (https://www.cgiar.org/funders/). Funding support was provided through the project “Development and implementation of a sustainable IPM and surveillance program for the invasive tomato leafminer, *Tuta absoluta* (Meyrick) in North and sub-Saharan Africa” financed by the German Federal Ministry for Economic Cooperation and Development (BMZ) and commissioned by the Deutsche Gesellschaft für Internationale Zusammenarbeit (GIZ) through the Fund International Agricultural Research (FIA) (Grant number: 81157481) to the International Centre of Insect Physiology and Ecology (*icipe*).

## Author Contribution

- Author 1, author 2, and author 3 conceived research.
- Author 1 conducted experiments.
- Author 3 and author 4 contributed material.
- Author 1 and 2 analysed data and conducted statistical analyses.
- Author 1, and author 3 wrote the manuscript.
- Author 3, and author 4 secured funding.
- All authors read and approved the manuscript.

## Data Availability Statement

Data are openly available in a public repository that issues datasets with DOIs. The data that support the findings of the life table and phenology modeling study of *Dolichogenidae gelechiidivoris* are available at https://data.cipotato.org/dataset.xhtml?persistentId=doi:10.21223/5VZE2H (Mujica and Carhuapoma, 2023a); and for *Tuta absoluta* at https://data.cipotato.org/dataset.xhtml?persistentId=doi:10.21223/BKLGMG (Mujica and Carhuapoma, 2023b).

