## Supplementary material for "Mapping of suitable release areas for the parasitoid *Dolichogenidea gelechiidivoris* (Marsh) for the classical biocontrol of *Tuta absoluta* (Meyrick) using temperature-dependent phenology models": Tables

**Table 1.** Mean development time (± SE), mortality, and model fitted to development time of immature *D. gelechiidivoris* life stages reared on *T. absoluta* at constant temperatures.

| Temperature (°C) | Egg-larvae | | | Pupae | | | Total | |
|---|---|---|---|---|---|---|---|---|
| | n | Median dev. time (days) | Mortality (%) | n | Median dev. time (days) | Mortality (%) | Median dev. time (days) | Mortality (%) |
| 10 | 32 | 60.225 (±8.92)a | 69.4 | - | - | 100 | - | 100 |
| 15 | 120 | 25.175 (±2.84)b | 18.4 | 98 | 27.635 (±1.195)a | 24.5 | 52.82 (±2.742)a | 38.3 |
| 20 | 240 | 15.753 (±1.937)bc | 32.5 | 120 | 11.883 (±0.649)b | 15.4 | 27.64 (±2.035)a | 26 |
| 25 | 240 | 10.968 (±1.535)c | 45 | 88 | 7.47 (±0.472)c | 16.7 | 18.44 (±1.68)b | 33.5 |
| 30 | 180 | 8.47 (±1.083)c | 56.7 | 78 | 5.773 (±0.443)c | 55.8 | 14.243 (±1.132)b | 80.5 |
| Model | | Loglogistic | | | Loglogistic | | | |
| Common slope | | 12. 992 | | | 13.329 | | | |
| P (>/z/) | | < 0.001 | | | < 0.001 | | | |
| AIC | | 1796.788 | | | 1038.447 | | | |

[a]Numbers in parenthesis are standard errors.

[b]Mean followed by different letters in the same columns are significantly different ($P < 0.05$; Tukey test). Egg-larvae: $F = 994.17$; df = 14,394; $P < 0.0001$. Pupae: $F = 771.76$; df = 12,265; $P < 0.0001$.

**Table 2.** Models and their parameters fitted to describe the median development rate (1 per d) for immature life stages of *D. gelechiidivoris* reared on *T. absoluta*

| Life stage | Models[a] | | Parameters | | *t*-value | $Df_{1,2}$ | *P* | Adj. $R^2$ |
|---|---|---|---|---|---|---|---|---|
| Egg-larvae | Janish-1 | $$r(T) = \frac{2}{D_{min} \times \left(e^{K\times(T-T_{opt})} + e^{-K\times(T-T_{opt})}\right)}$$ | $D_{min}$ | 8.577 (±0.721)[b] | 11.89 | 2,12 | < 0.0001 | 0.947 |
| | | | $T_{opt}$ | 29.632 (±1.601) | 18.51 | | < 0.0001 | |
| | | | K | 0.129 (±0.012) | 10.93 | | < 0.0001 | |
| Pupae | Janish-1 | $$r(T) = \frac{2}{D_{min} \times \left(e^{K\times(T-T_{opt})} + e^{-K\times(T-T_{opt})}\right)}$$ | $D_{min}$ | 5.817 (±0.276) | 21.07 | 2,10 | < 0.0001 | 0.976 |
| | | | $T_{opt}$ | 29.288 (±0.837) | 34.98 | | < 0.0001 | |
| | | | K | 0.153 (± 0.011) | 14.1 | | < 0.0001 | |

[a]Janish-1: where r(T) is the development rate at temperature T (°C), $T_{opt}$ the temperature at which the development rate is at maximum, and $D_{min}$ and K are fitted constants.

[b]Numbers in parenthesis are standard errors.

**Table 3.** Model and their parameters fitted to describe mortality rate for immature life stages of *D. gelechiidivoris* reared on *T. absoluta*

| Life stage | Model | | Parameters | | *t*-value | $Df_{1,2}$ | P |
|---|---|---|---|---|---|---|---|
| | | | a | 0.015 (±0.004)[b] | 3.329 | | 0.010 |
| Egg-larvae | | $m(T) = \frac{e^{(a \times T^2 + b \times T + c)}}{e^{(a \times T^2 + b \times T + c)} + 1}$ | b | -0.603 (±0.186) | -3.244 | 2,8 | 0.012 |
| | | | c | 5.023 (±1.731) | 2.902 | | 0.019 |
| | Quadratic[a] | | | | | | |
| | | | a | 0.026 (±0.006) | 3.845 | | 0.003 |
| Pupae | | $m(T) = \frac{e^{(a \times T^2 + b \times T + c)}}{e^{(a \times T^2 + b \times T + c)} + 1}$ | b | -0.989 (±0.214) | -4.62 | 2,9 | 0.001 |
| | | | c | 6.783 (±1.917) | 3.538 | | 0.006 |

[a]Quadratic: where *m(T)* is the rate of mortality at temperature T (°C), *a*,*b* and *c* are the fitted parameters.

[b]Numbers in parenthesis are standard errors.

.

**Table 4** Median survival time, median oviposition time, mean fecundity, and models fitted to describe these development parameters of *D. gelechiidivoris* reared on *T. absoluta* at constant temperatures

| Temperature (°C) | Longevity (days) | | Median oviposition time (days) | Mean fecundity (eggs/female) |
|---|---|---|---|---|
| | Female | Male | | |
| 10 | 36.36 (±3.52) [a]a[b] | 26.72 (±2.05) ab | 16.655 (±3.276) ab | 23.165 (±1.632) a |
| 15 | 23.13 (±3.16) b | 29.3 (±3.15) a | 9.796 (±2.306) abc | 50.52 (±1.94) b |
| 20 | 15.5 (±2.14) b | 20.1 (±2.16) ab | 6.326 (±1.427) abcd | 74.355 (±3.124) c |
| 25 | 8.83 (±1.21) c | 8.64 (±0.93) c | 3.445 (±0.811) cd | 55.043 (±2.573) b |
| 30 | 8.9 (±1.22) c | 8.59 (±0.93) c | 3.579 (±0.982) bcd | 23.918 (±1.29) a |
| 35 | 3.6 (±0.5) d | 3.56 (±0.39) d | 1.622 (±0.597) d | 12.452 (±0.848) d |
| Model | Weibull | Weibull | Weibull | Taylor |
| Common slope | 2.5908 | 2.8916 | 1.31658 | |
| P (>/z/) | < 0.001 | < 0.001 | < 0.001 | |
| AIC | 1930.64 | 1846.143 | | -8.5096 |

[a]Numbers in parenthesis are standard errors.

[b]Means followed by different letters in the same columns are significantly different ($P < 0.05$; Tukey test). Female: $F = 420.63$; df = 5, 300; $P < 0.0001$. Male: $F = 441.57$; df = 5, 300; $P < 0.0001$. Oviposition: $F = 9591.94$; df = 5, 22531; $P < 0.0001$).

**Table 5.** Estimated parameters of the non-linear models fitted to describe the relationship between temperature and adult senescence rates, oviposition rate, and fecundity for *D. gelechiidivoris* reared on *T. absoluta*

| Response variable | Model | | Parameters | | *t*-value | $df_{1,2}$ | P |
|---|---|---|---|---|---|---|---|
| | | | a | 3e-04 ($\pm$0.0396)[c] | 1.056 | | 0.147 |
| Female senescence rate | Quadratic[a] | $r(T) = \frac{1}{e^{(a \times T^2 + b \times T + c)}}$ | b | -0.004 ($\pm$0.0049) | -0.808 | 2,3 | 0.477 |
| | | | c | 0.042 ($\pm$0.0013) | 1.943 | | 0.368 |
| | | | a | 0.112 ($\pm$0.0413) | 3.435 | | 0.041 |
| Male senescence rate | Quadratic | $r(T) = \frac{1}{e^{(a \times T^2 + b \times T + c)}}$ | b | -0.012 ($\pm$0.005) | -2.472 | 2,3 | 0.089 |
| | | | c | 4e-04 ($\pm$0.0001) | 2.705 | | 0.073 |
| | | | a | 0.065 ($\pm$0.081) | 1.932 | | 0.148 |
| Oviposition rate | Quadratic | $r(T) = \frac{1}{e^{(a \times T^2 + b \times T + c)}}$ | b | -0.005 ($\pm$0.0101) | -0.566 | 2,3 | 0.611 |
| | | | c | 5e-04 ($\pm$0.0003) | 0.799 | | 0.482 |
| | | | $r_m$ | -3.263 (+0.059) | -55.11 | | 0.999 |
| Fecundity | Taylor[b] | $n(T) = 1 - r_m \times e^{\frac{-1}{2}\left(\frac{T_{opt} - x}{T_{roh}}\right)^2}$ | $T_{op}$ | 20.35 ($\pm$0.274) | 74.23 | 2,3 | <0.001 |
| | | | $T_{roh}$ | 11.414 ($\pm$0.393) | 29.07 | | <0.001 |

[a]Quadratic: r(T) is the senescence rate at temperature T (°C), where a, b and c are equation parameters. [b]Taylor: n(T) represents the fecundity function at temperature T (°C) and $r_m$, $T_{opt}$ and $T_{roh}$ are parameters of the equation. [c]Numbers in parenthesis are standard errors.

**Table 6.** Statistic life table summary from simulated and observed life table parameters, development time, and mortality of *D. gelechiidivoris* reared on *T. absoluta* at fluctuating temperatures

| | Simulated | | Observed | *P* |
|---|---|---|---|---|
| **Life table parameters** | | | | |
| Intrinsic rate of increase (*r*) | 0.014 | (± 0.047) | 0.044 | 0.2226 |
| Mean generation time (*T*) | 39.473 | (± 7.3) | 40.286 | 0.7155 |
| Finite rate of increase (λ) | 1.014 | (± 0.048) | 1.045 | 0.2199 |
| **Development time (days)** | | | | |
| Egg-larvae | 19.441 | (± 0.46) | 21.784 | 0.0021 |
| Pupa | 9.37 | (± 0.415) | 12.361 | 0.001 |
| **Mortality (%)** | | | | |
| Egg-larvae | 0.525 | (± 0.231) | 0.15 | 0.0166 |
| Pupae | 0.241 | (± 0.126) | 0.294 | 0.1184 |
